# Building the meiotic bouquet: A nuclear polarity pathway in Arabidopsis

**DOI:** 10.64898/2026.05.27.728084

**Authors:** Michaela Kubova, Martin A. Lysak, Terezie Mandakova

## Abstract

Meiotic chromosome pairing requires the establishment of a polarized nuclear architecture, but how this polarity emerges from the interplay of telomeres, the nucleolus, and the nuclear envelope remains incompletely understood in plants. Using stage-resolved three-dimensional imaging of intact *Arabidopsis thaliana* male meiocytes, we reconstruct the early meiotic sequence that links the premeiotic chromosome arrangement to telomere bouquet formation. We show that telomeres enter meiosis positioned around the nucleolus and, at the onset of leptotene, undergo a coordinated and directionally biased departure from the nucleolar surface. This transition establishes asymmetric early contacts with discrete SUN1/2-enriched domains of the nuclear envelope, well before visible clustering occurs, revealing that the envelope is pre-patterned into regions permissive for telomere attachment. As zygotene progresses, these early contacts consolidate into a nucleolus-associated bouquet, whereas the nucleolar organizer regions (NORs) maintain a stable position and remain unsynapsed. Centromeres reorganize in parallel and progressively shift away from the developing telomere domain, giving rise to the classical centromere–telomere opposition. Rare nuclei exhibiting bouquet formation opposite the nucleolus correspond to alternative SUN1/2 microdomain topologies, indicating that nuclear-envelope patterning, rather than telomere behavior, ultimately determines the polarity state. Together, these findings provide a mechanistic framework in which NOR stability, SUN-patterned membrane domains, and telomere migration act sequentially to generate a polarized meiotic nucleus in Arabidopsis.

## INTRODUCTION

Meiotic chromosome segregation requires chromosomes to move, interact and reorganize within the confined nuclear space, making early prophase I one of the most architecturally dynamic phases of the cell cycle. During leptotene and zygotene, chromosomes undergo extensive spatial reconfiguration, including the establishment of nuclear polarity, redistribution of chromosome ends, and formation of the telomere bouquet - a conserved arrangement in which telomeres cluster at a restricted region of the nuclear envelope (NE) to promote homolog recognition (Zickler & Kleckner, 1998; Tomita & Cooper, 2007). These transitions are driven by cytoskeletal forces transmitted to chromosome ends through LINC-complex proteins embedded in the NE, and together they create the structural framework that precedes and facilitates homolog pairing (Sheehan & Pawlowski, 2009; Fernández-Álvarez, 2023). Although bouquet formation is conserved across eukaryotes, its timing, geometry and nuclear context vary between systems, underscoring the need to understand how plant meiocytes, with their large nuclei and prominent nucleoli, establish this early meiotic polarity.

Although telomere clustering is broadly conserved, the spatial context in which bouquet formation occurs differs substantially across eukaryotes. In budding and fission yeast, telomeres attach to a defined microdomain of the nuclear envelope near the spindle pole body, driving rapid chromosome oscillations that facilitate homolog encounters (Tomita & Cooper, 2007; Fernández-Álvarez, 2023). In vertebrates such as zebrafish, bouquet formation coincides with vigorous telomere-led motions and localized initiation of synapsis events (Blokhina et al., 2019). Plants exhibit the same overall choreography - telomere migration to the NE during leptotene, increasing nuclear polarity, and zygotene bouquet formation - but do so within much larger nuclei dominated by a prominent nucleolus, which undergoes its own characteristic repositioning (Cowan et al., 2001; Sheehan & Pawlowski, 2009). As a consequence, plant meiocytes provide a particularly clear system in which to study how telomere migration, nucleolar displacement, and NE attachment collectively shape early meiotic architecture.

In *Arabidopsis thaliana* (Arabidopsis), early prophase I is characterized by a highly ordered sequence of spatial transitions that has been described in both fixed and live meiocytes. Telomeres, which frequently occupy positions adjacent to the nucleolus during premeiotic interphase, begin to migrate outward as leptotene initiates and form numerous individual foci along the inner NE before any visible clustering arises (Sheehan & Pawlowski, 2009; Hurel et al., 2018). This relocalization depends on intact chromosome-NE interactions mediated by the LINC complex: the inner nuclear membrane SUN-domain proteins SUN1 and SUN2 are essential for telomere tethering and rapid chromosome motions, and their disruption leads to loss of polarized telomere organization and defects in synapsis (Varas et al., 2015; Cromer et al., 2024). In parallel, the nucleolus undergoes a marked repositioning toward the nuclear periphery, creating an early nuclear asymmetry that is a hallmark of the leptotene-zygotene transition in plants. As telomere polarization becomes established, centromeres - which tend to accumulate near the nucleolar side during early stages - progressively redistribute, eventually giving rise the characteristic telomere-centromere opposition observed in plant meiocytes (Cowan et al., 2001; Cromer et al., 2024). Together, these observations outline a framework in which telomere migration, nucleolar repositioning, and SUN-dependent nuclear-envelope interactions cooperate to establish meiotic nuclear polarity in Arabidopsis.

Despite these detailed cytological descriptions, several critical aspects of early meiotic polarity in Arabidopsis remain unresolved. First, the spatiotemporal sequence by which telomeres transition from their premeiotic nucleolar-proximal positions to stable attachment sites on the nuclear envelope - and how this transition gives rise to a polarized bouquet - has not been reconstructed in intact, stage-resolved 3D meiocytes. Second, although telomere–NE tethering depends on SUN1 and SUN2, the spatial organization of SUN-enriched membrane domains around the nuclear periphery, and their potential role in defining where bouquets form, is still poorly understood. Third, how the bouquet relates to major nuclear landmarks, including the nucleolus, nucleolar organizing regions (NORs), and the characteristic centromere–telomere opposition, has not been integrated into a single architectural framework. Finally, while recent studies suggest non-canonical roles for telomeres and NE remodeling in organizing meiotic chromosomes, the mechanistic logic linking telomere migration, nucleolar relocation, and SUN-mediated anchoring remains unclear (Zeng et al., 2018; Fernández-Álvarez, 2023). Together, these gaps highlight the need for a unified, spatially explicit model of how meiotic nuclear polarity emerges in plants.

To address these open questions, we performed a stage-resolved three-dimensional analysis of intact Arabidopsis male meiocytes, combining immunolabeling of key meiotic axis and NE components with FISH detection of telomeres, centromeres and 35S rDNA. This integrated cytological framework allowed us to follow, within the same staged nuclei, the coordinated behavior of chromosome ends, NORs, and SUN1/2-enriched membrane domains throughout leptotene and zygotene. By reconstructing the sequence of events from the premeiotic nucleus to bouquet formation, we uncover how telomere migration, nucleolar repositioning, and patterned nuclear-envelope attachment jointly establish meiotic nuclear polarity. This approach not only refines the temporal order of early meiotic events but also reveals previously unrecognized intermediate states and structural constraints that organize bouquet placement in Arabidopsis.

## RESULTS

### The nucleolus relocates from a central to a peripheral nuclear position during early prophase I

During premeiotic interphase (**Fig. 1A–D**), the nucleolus occupies a central position within the meiocyte nucleus. Fibrillarin immunolabeling and 35S rDNA FISH signals form a single compact nucleolar domain, with the rDNA arrays positioned immediately adjacent to and partially overlapping with the fibrillarin-positive region. As cells enter leptotene (**Fig. 1E–H**), the nucleolus begins a reproducible shift away from the nuclear center and adopts an eccentric position that remains tightly associated with the 35S rDNA signals. This displacement becomes more pronounced during zygotene (**Fig. 1I–L**), at which point the nucleolus consistently occupies a peripheral site at one side of the nucleus. By pachytene (**Fig. 1M–P**), the nucleolus retains this peripheral, crescent-shaped configuration, forming a robust spatial landmark in an otherwise highly reorganized chromatin environment. The staged and directional nature of this relocation is fully consistent across all nuclei examined and provides an unambiguous reference point for ordering early meiotic prophase I events.

**Figure 1.**
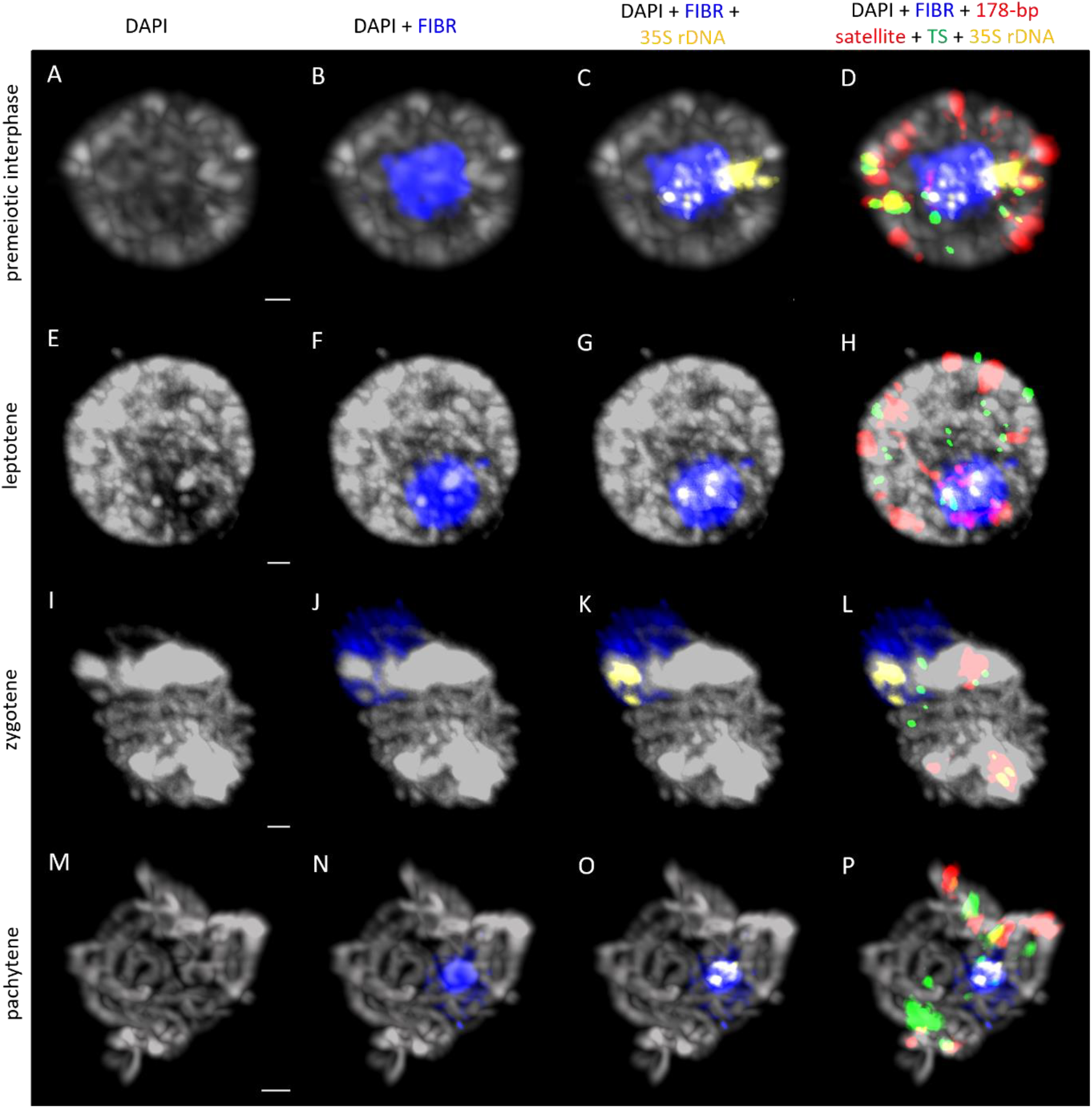
Identification of the nucleolus in Arabidopsis meiocytes throughout premeiotic interphase, zygotene, and pachytene. 3D immunodetection of nucleoli using anti-fibrillarin antibody (FIBR, blue) followed by FISH of the 35S rDNA probe (yellow); centromeres were visualized using the 178-bp centromeric satellite probe (red) and telomeres using Arabidopsis-like telomeric repeat (green). Chromosomes were counterstained by DAPI (grey). Scale bars: 1 µm (A–L), 2 µm (M–P).

### ASY1 and ZYP1 signal dynamics delineate the progression of meiotic chromosome pairing

To resolve the sequence of chromosome-pairing events during early prophase I, we performed immunolabeling of the axis protein ASY1 and the synaptonemal complex (SC) central-element protein ZYP1 in polyacrylamide-embedded meiocytes (**Fig. 2**). During premeiotic interphase (**Fig. 2A**), ASY1 appears as scattered puncta on decondensed chromatin, while ZYP1 is undetectable, marking a meiotic G2 stage. At leptotene (**Fig. 2B**), ASY1 forms continuous linear axes along each chromosome, coincident with chromatin condensation into axial elements. At early zygotene (**Fig. 2C**), ZYP1 emerges as discrete punctate foci at multiple sites along ASY1-marked axes, corresponding to synapsis initiation centers. ASY1 remains present on unsynapsed regions, enabling simultaneous visualization of both synapsed and unsynapsed chromosome segments. As zygotene progresses (**Fig. 2D–M**), ZYP1 extends into longer stretches marking growing SC segments, while ASY1 signal diminishes from regions undergoing synapsis, consistent with its displacement from fully synapsed axial elements.

**Figure 2.**
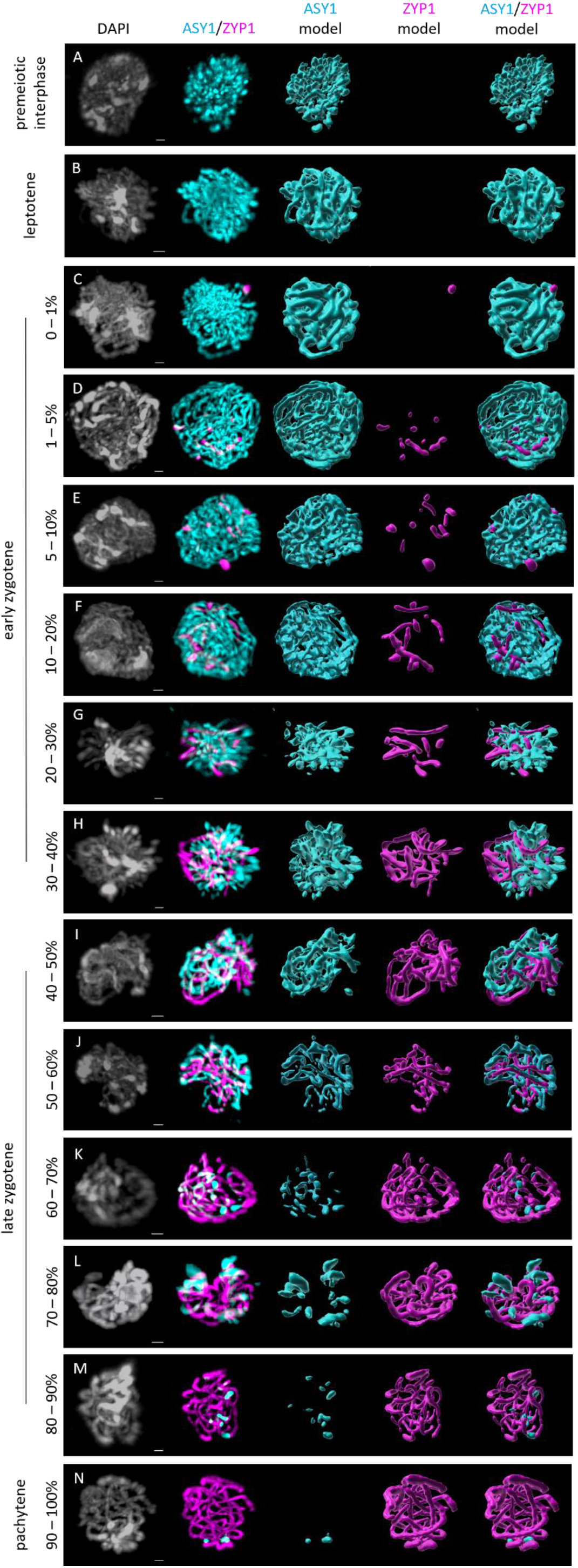
Dynamics of ASY1 and ZYP1 localization during meiotic prophase I in Arabidopsis male meiocytes. Immunolocalization of the synaptonemal complex proteins ASY1 (cyan) and ZYP1 (magenta) in polyacrylamide-embedded meiocytes. (A) During premeiotic interphase, ASY1 appears as punctate foci on decondensed chromatin, while ZYP1 is absent. (B) At leptotene, chromatin condenses into axial elements marked by continuous ASY1 signal. (C, D) The initiation of synapsis during early zygotene is marked by the emergence of ZYP1 foci. (E–M) As zygotene progresses, ZYP1 extends along chromosomes and ASY1 is progressively depleted from synapsed regions. (N) By pachytene, full synapsis is indicated by continuous ZYP1 signals along entire bivalents, with ASY1 foci persistent only at NORs (see Fig. 3). Percentage values indicate the relative abundance of ZYP1 signals quantified by the Surface and Vantage functions in IMARIS. Scale bars, 1 µm.

By pachytene (**Fig. 2N**), ZYP1 forms continuous signals along all bivalents, indicating complete synapsis throughout the genome except at the NORs (**Fig. 3**). ASY1 persists only at this single, discrete locus. Quantification of ZYP1 abundance using the IMARIS Surface and Vantage tools (percentage values in **Fig. 2**) provides a robust, quantitative staging framework that supports fine-scale temporal ordering of meiotic nuclei across leptotene and zygotene.

**Figure 3.**
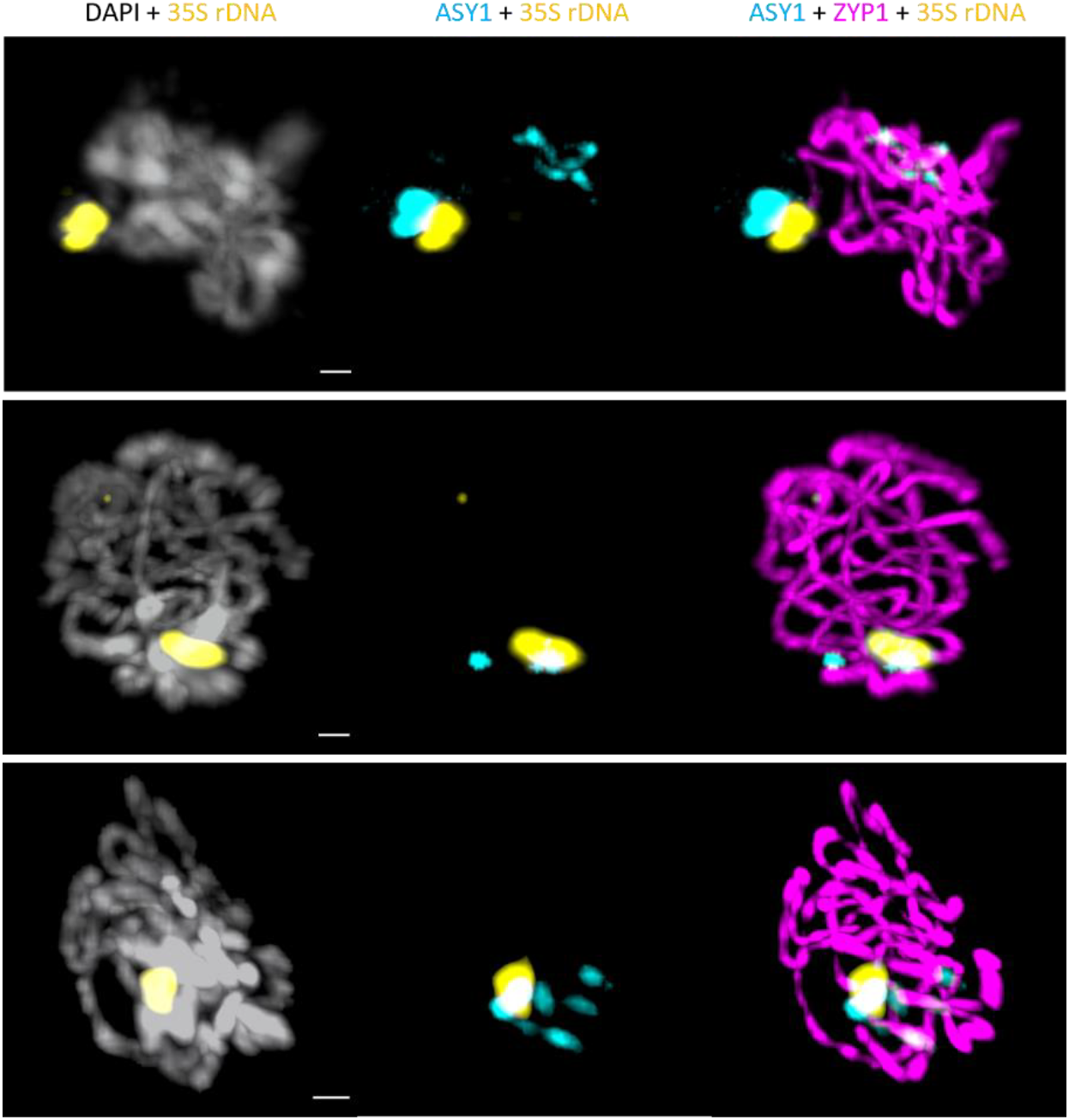
Persistent ASY1 signal colocalizes with the 35S rDNA in pachytene. Immunolocalization of ASY1 (cyan) at a clustered NORs detected with the 35S rDNA probe (yellow). Prophase I staging was determined based on continuous ZYP1 signals (magenta) in fully synapsed bivalents (see also Fig. 2). Meiocytes were counterstained with DAPI (grey). Scale bars, 1 µm.

### The NORs remain unsynapsed and retains ASY1 during early pachytene

To identify the chromosomal domain that retains ASY1 after genome-wide synapsis, we combined ASY1 immunolabeling with FISH detection of the 35S rDNA arrays in pachytene meiocytes (**Fig. 3**). In all nuclei examined, a single, bright ASY1 focus persisted at a position that precisely colocalized with the 35S rDNA signals marking the NORs. Continuous ZYP1 labeling along all other chromosome arms confirmed complete synapsis across the genome, whereas ZYP1 was consistently absent at the NORs. NOR-associated chromatin has been shown to exhibit strongly reduced DSB formation and minimal engagement of homologous recombination during meiosis (Sims et al., 2019; Sims et al., 2021), and the lack of ZYP1 together with the persistent ASY1 signal observed here reflects this distinctive behavior. The ASY1-positive NOR therefore provides a reliable cytological marker for identifying fully synapsed pachytene nuclei.

### Spatial reorganization of telomeres, centromeres, and NORs during early meiotic prophase I

Because the NE was not immunolabeled in these experiments, all positional descriptions in this section refer to the nuclear periphery, rather than direct NE attachment. NE association is addressed separately in the subsequent section using SUN1/2 immunolabeling. To examine the spatial dynamics of chromosome landmarks during early meiotic prophase I, we performed 3D FISH using probes targeting telomeric repeats, the 178-bp pericentromeric satellite, and the 35S rDNA arrays, and staged meiocytes based on ASY1 and ZYP1 signal patterns (**Fig. 2, Fig. 4**). During premeiotic interphase (**Fig. 4A**; **Video 1**), telomeres were predominantly positioned around the nucleolus and frequently appeared in close proximity to the 35S rDNA signals; in 94% of nuclei (n = 18), telomeres and both NOR loci were associated with a single nucleolus. Pericentromeric heterochromatin localized to peripheral nuclear regions. A prominent interstitial telomeric repeat (ITR) was consistently detected at the (peri)centromeric region of chromosome 1 and remained visible at all subsequent meiotic stages.

**Figure 4.**
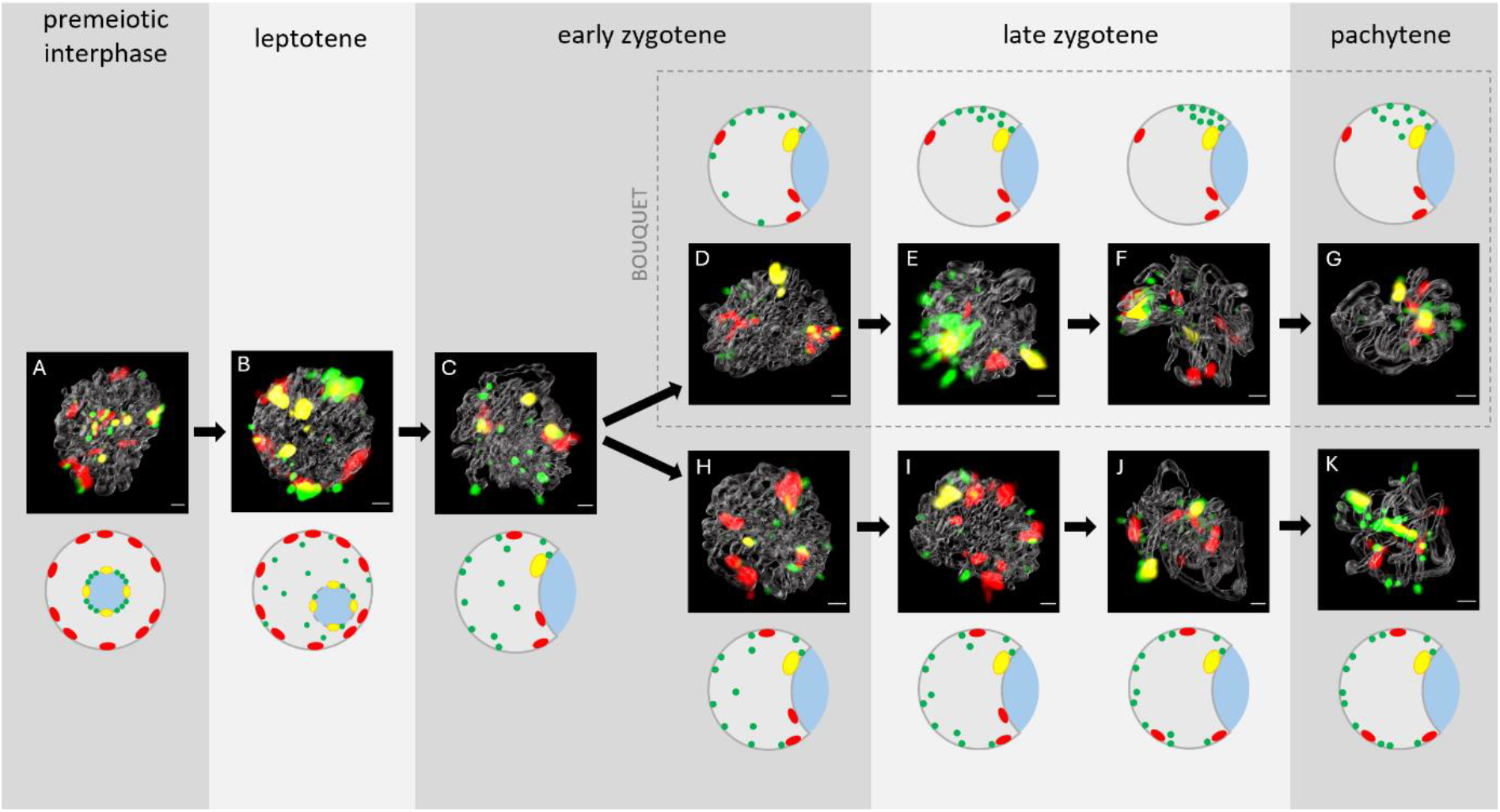
Spatiotemporal chromosome dynamics in intact Arabidopsis male meiocytes during G2 premeiotic interphase and early meiotic prophase I. Meiotic stages were identified using ASY1 and ZYP1 immunolabeling (grey; see also Fig. 2). Telomeric signals (green), 178-bp centromeric satellite signals (red), and 35S rDNA loci (yellow) were visualized by FISH. Schematic diagrams illustrate the repositioning of telomeres, centromeres, and 35S rDNA loci during meiotic progression. (A) premeiotic interphase, (B) leptotene, and (C) early zygotene, representing the initial phase of large-scale nuclear reorganization. (D–G) Zygotene to pachytene stages characterized by the establishment of telomere clustering at the nuclear periphery. (H–K) Zygotene and pachytene nuclei illustrate the prevalent patterns of chromosome movement associated with homolog pairing and synapsis. In all diagrams, cobalt blue denotes the nucleolus. Scale bars: 1 µm.

As cells entered leptotene (**Fig. 4B**; **Video 2**), telomeres initiated a coordinated displacement away from the nucleolus toward the nuclear periphery, where they formed peripheral or sub-peripheral distributions. During this transition, telomeric foci frequently interlaced with adjacent pericentromeric regions. Centromeres generally maintained peripheral positions, positioned opposite or lateral to the nucleolus. The 35S rDNA loci remained associated with the nucleolus in all leptotene nuclei examined (n = 17).

During early and late zygotene (**Fig. 4C–F, H–J**; **Videos 3–6, 8–10**), telomeres were detected at the nuclear periphery in 76% of nuclei (n = 63), forming a broad peripheral arrangement. Approximately 20% of nuclei displayed partial interior telomere dispersion, representing transitional states during relocalization. Centromeres exhibited two predominant configurations: clustering near the nucleolus together with 35S rDNA loci (59% of nuclei) or dispersion within the nuclear interior (32%). Across leptotene and zygotene, 35S rDNA loci remained nucleolus-associated in 97% of nuclei, providing a consistent positional landmark within the reorganizing nucleus.

At pachytene (**Fig. 4G, K**; **Videos 7, 11**), dense interweaving of fully synapsed chromosomal axes impeded resolution of individual landmark positions. Telomeres remained detectable at the nuclear periphery, consistent with their earlier distributions. The positions of centromeres and NORs relative to the nuclear periphery could not be reliably determined due to the increased axial density of fully synapsed bivalents.

### Telomere bouquet formation during the leptotene–zygotene transition

To quantify telomere bouquet formation, we analyzed telomeric, centromeric, and 35S rDNA signals in staged meiocytes using 3D FISH (**Fig. 5**). A bouquet was defined as a compact cluster of telomeric foci located within a restricted region of the nuclear periphery. Bouquet structures were identified in 32% of all meiocytes analyzed (41/128). In all bouquet-positive nuclei, telomeres localized to the nuclear periphery; occasional detection of individual interior telomeric signals likely reflects transitional states captured during fixation. Telomeres associated with NORs (chromosomes 2 and 4) and the ITR on chromosome 1 were excluded from bouquet scoring.

**Figure 5.**
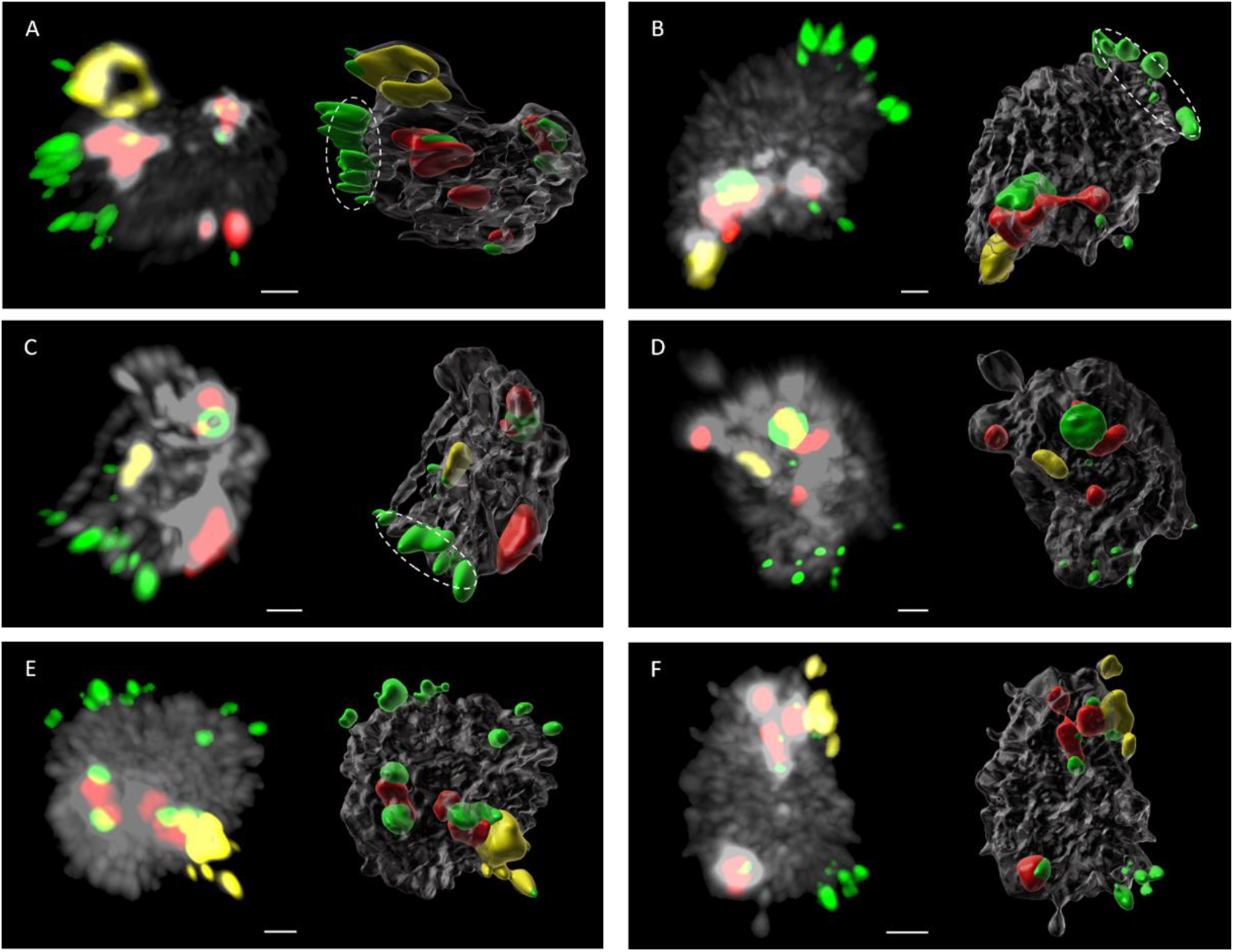
Telomere bouquet formation during early meiotic prophase I. (A) Late leptotene/zygotene meiocyte displaying a telomere bouquet (dashed ellipse) formed by a tight cluster of telomeric signals (green) at the NE adjacent to the nucleolus, identified by the 35S rDNA signal (yellow). Centromeres were visualized with the 178-bp centromeric satellite probe (red). (B) Alternative bouquet configuration in a late leptotene/zygotene meiocyte, with telomeres clustered at the NE on the hemisphere opposite the nucleolus. (C) Pachytene-stage meiocyte showing tightly paired telomeric signals aligned along the NE. (D–F) Representative examples of the three centromere-positioning patterns observed in bouquet-positive nuclei: (D) centromeres clustered near the 35S rDNA locus, (E) centromeres dispersed throughout the nuclear interior, and (F) centromeres distributed along the NE, with most located close to the 35S rDNA signal. For each panel pair, the left image shows the DAPI-stained meiocyte together with FISH signals, and the right image shows the corresponding Imaris-generated 3D reconstruction. Meiocytes were counterstained with DAPI (grey). Scale bars: 1 µm (B, D); 2 µm (A, C, E, F).

Bouquets were most frequent during late leptotene and zygotene: 39 of 41 bouquet-positive nuclei corresponded to these stages. The number of telomeric signals within a bouquet ranged from 5 to 19 (mean 10.7). Two spatial configurations were observed. In 88% of nuclei (n = 36), bouquets were positioned at the nuclear periphery adjacent to the nucleolus (**Fig. 5A**). In 12% (n = 5), bouquets were located at the nuclear periphery opposite the nucleolus (**Fig. 5B**).

Centromere positioning in bouquet-positive leptotene/zygotene nuclei showed three common patterns (**Fig. 5D–F**): clustering near the nucleolus (77%, 30/39 nuclei), dispersion within the nuclear interior (13%, 5/39), or peripheral distribution with most signals near the nucleolus (10%, 4/39). At pachytene, bouquet-like arrangements were rare (2/41 nuclei), with paired telomeric signals aligned along the nuclear periphery (**Fig. 5C**), displaying on average 10 telomeric and 3.5 centromeric foci. Due to dense axial organization, centromere and NOR positions could not be reliably resolved in most pachytene nuclei.

### SUN1/2-labeled nuclear envelope domains reveal telomere association with discrete membrane regions

To directly visualize telomere contacts with the NE, we performed immunolabeling with an anti-SUN1/2 antibody, which marks the inner nuclear membrane, followed by FISH detection of telomeres, centromeres, and 35S rDNA (**Fig. 6**). In contrast to the previous section, where only the nuclear periphery could be assessed, this approach enabled direct determination of NE association.

**Figure 6.**
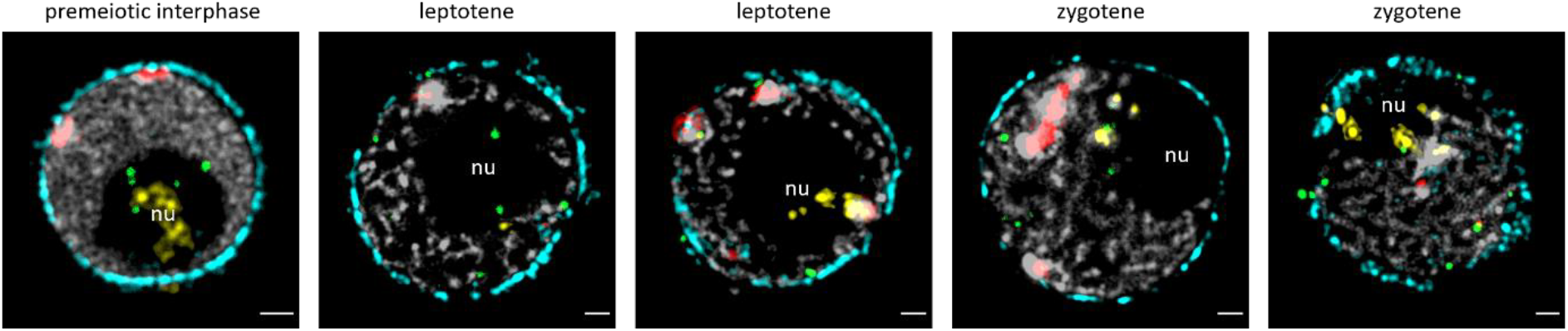
Nuclear envelope organization and telomere positioning during early meiotic prophase I. 3D immunolocalization of the NE using an anti-SUN1/2 antibody (cyan), combined with FISH detection of telomeric repeats (green), the 178-bp centromeric satellite probe (red), and 35S rDNA loci (yellow) in intact Arabidopsis meiocytes. During premeiotic interphase, telomeres localize predominantly at the nucleolar periphery (nu), with occasional foci detected at the NE. In leptotene, telomeres are observed both at the nucleolar membrane and at SUN-labeled NE regions, whereas in zygotene they localize predominantly to SUN-enriched domains of the NE. All images represent projections of selected z-sections acquired by epifluorescence microscopy followed by deconvolution. Meiocytes were counterstained with DAPI (grey). Scale bars: 1 µm.

The anti-SUN1/2 antibody consistently labeled the NE, revealing that SUN1/2 signals were not uniformly distributed, but formed discrete microdomains along the envelope. During premeiotic interphase, telomeres localized predominantly around the nucleolus, consistent with their positions described above, with only occasional telomeric foci contacting the NE (**Fig. 6A**). In leptotene (**Fig. 6B**), telomeres were detected both at the nucleolar surface and at SUN1/2-labeled NE regions, indicating the onset of NE association. During zygotene (**Fig. 6C**), telomeres localized predominantly to SUN1/2-positive domains of the NE, forming multiple perinuclear foci distributed along SUN-enriched membrane regions.

In all staged nuclei examined, telomeres positioned at the NE overlapped spatially with SUN1/2 microdomains rather than with SUN-poor membrane regions. Quantification showed that in 81% of leptotene nuclei (n = 21) and 87% of zygotene nuclei (n = 31), SUN1/2 signal intensity was enriched at the nuclear hemisphere containing the majority of telomeric foci. This spatial correspondence indicates that telomere–NE contacts in Arabidopsis occur preferentially at discrete SUN1/2-enriched membrane regions. The positions of centromeres and 35S rDNA loci in SUN-labeled nuclei matched the distributions described in previous sections.

## DISCUSSION

### Bouquet formation during early meiosis proceeds through intermediate states

Our staged 3D analysis confirms key aspects of early meiotic chromosome behavior previously described in Arabidopsis, while extending them by resolving intermediate configurations not previously visualized in intact meiocytes. Consistent with classical cytology and live imaging (Armstrong et al., 2001; Prusicki et al., 2019), telomeres began meiosis in close proximity to the nucleolus and then redistributed toward the nuclear periphery as cells entered leptotene, in agreement with earlier descriptions of plant chromosome-end movement (Sheehan & Pawlowski, 2009; Hurel et al., 2018). By capturing nuclei across a finely staged leptotene–zygotene continuum, our dataset clarifies that this redistribution occurs through gradual, spatially biased transitions rather than through an abrupt relocalization. Bouquet formation in late leptotene and zygotene consisted of its established developmental timing (Armstrong et al., 2001) and exhibited the expected nucleolar-side bias. The identification of a small subset of nuclei with bouquets positioned on the opposite nuclear hemisphere, a feature not documented in earlier Arabidopsis studies, suggests that bouquet polarity is consistent, but not invariant. Collectively, these observations provide a more continuous and spatially resolved view of early chromosome-end dynamics and establish intermediate positional states that precede bouquet formation.

### NORs constitute a stable nuclear landmark in an otherwise dynamic meiotic nucleus

NORs exhibited a striking positional and structural stability throughout early prophase I, maintaining close association with the nucleolus from premeiotic interphase through pachytene. This behavior aligns with previous studies showing that rDNA arrays form a recombination-refractory chromatin domain that neither undergoes meiotic double-strand break formation nor engages in canonical homologous synapsis (Sims et al., 2019; Sims et al., 2021). In our data, NOR stability contrasted with the extensive repositioning of telomeres and centromeres that accompanies nuclear polarization during leptotene and zygotene. By retaining both their position and unsynapsed state even as other chromosomal regions reorganize, NORs serve as a robust spatial reference within the transforming nucleus. The combination of NOR immobility and the high spatial resolution of intact 3D meiocytes, the stability allowed chromosome-end trajectories to be anchored relative to a fixed region, thereby clarifying how early nuclear polarity unfolds around a structurally stable chromatin domain.

### Bouquet position reflects the distribution of SUN1/2-enriched nuclear-envelope domains

The nuclear envelope of Arabidopsis meiocytes displayed discrete SUN1/2-enriched domains rather than a uniform distribution of SUN proteins, consistent with prior analyses using structured illumination and SUN-deficient mutants (Varas et al., 2015; Cromer et al., 2024). By integrating SUN1/2 immunolabeling with telomere and NOR detection in intact staged nuclei, our dataset shows how these membrane microdomains relate spatially to early chromosome-end behavior. During leptotene and zygotene, telomeres associated with the nuclear envelope predominantly overlapped with SUN-enriched regions, while SUN-poor areas rarely exhibited telomere contacts. This correspondence indicates that telomere tethering occurs preferentially at SUN1/2-rich domains rather than across the nuclear envelope uniformly. A minority of nuclei exhibited SUN1/2 enrichment on the opposite hemisphere, and in these nuclei telomere clustering occurred correspondingly on that opposite side.

These alternative bouquet positions did not correspond to specific meiotic substages and likely reflect natural variation in SUN domain topology. Together, these findings suggest that bouquet polarity reflects the distribution of SUN1/2 microdomains, and that the nucleolar-side bias arises because SUN enrichment most frequently develops next to the nucleolus.

### Functional implications and a working model for early meiotic nuclear polarity

The spatial patterns uncovered in our staged 3D analysis provide a framework for understanding how early meiotic polarity emerges and how it may influence the nuclear environment in which homologous chromosomes first encounter one another. In multiple organisms, bouquet formation has been proposed to facilitate early homolog interactions by concentrating chromosome ends in a circumscribed nuclear region and enabling coordinated chromosome movements (Zickler & Kleckner, 1998; Tomita & Cooper, 2007; Blokhina et al., 2019). While our data do not directly assess pairing initiation, the reproducible nucleolar-side bias of bouquet placement - and its correspondence to SUN1/2-enriched nuclear-envelope domains - indicates that chromosome ends are consistently brought into proximity within a defined nuclear sector during leptotene and zygotene. The conjunction of bouquet positioning and NOR stability thus establishes a persistent polarity axis along which early meiotic chromosome movements occur.

Integrating chromosome-end, NOR and SUN1/2 positions across staged nuclei enables us to propose a working model for the coordinated establishment of early meiotic polarity (**Fig. 7**). In this model, the NOR-associated nucleolus functions as a stable reference point within a reorganizing nucleus. As leptotene initiates, telomeres move away from the nucleolar periphery toward the nuclear envelope through progressively polarized intermediate states. Discrete SUN1/2-enriched membrane domains - frequently positioned adjacent to the NOR - provide preferred sites for telomere association at the nuclear surface. Bouquet formation during zygotene arises from the consolidation of these SUN-associated telomere positions, producing the dominant nucleolar-side bouquet configuration. In rare cases where SUN1/2-enriched regions localize to the opposite hemisphere, telomeres cluster on that side accordingly, indicating that bouquet position follows SUN-domain topology rather than being strictly determined by NOR proximity.

**Figure 7.**
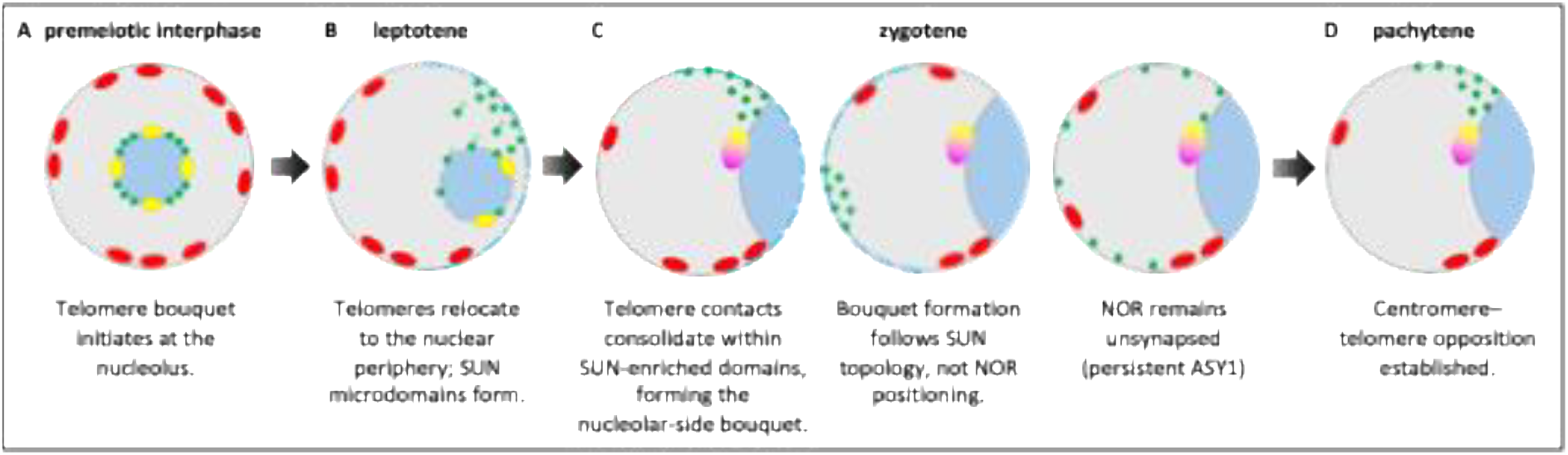
Model of early meiotic nuclear polarity in Arabidopsis. The model summarizes the sequence of spatial events observed during early meiotic prophase I, integrating telomere behavior (green), positions of 35S rDNA/NORs (yellow), the 178-bp centromeric satellite probe (red), the fibrillarin-marked nucleolus (cobalt blue), anti-SUN1/2 labeling (cyan), and the persistent single ASY1 focus (yellow/magenta). (A) During premeiotic interphase, telomeres are predominantly positioned at the periphery of the NOR-associated nucleolus, whereas centromeres occupy peripheral regions of the nucleus. At this stage, the NOR–nucleolus complex provides a stable spatial landmark. (B) At the onset of leptotene, telomeres disengage from the nucleolar periphery and progressively approach the nuclear envelope (NE) through increasingly asymmetric intermediate configurations. Initial contacts frequently occur at SUN1/2-enriched domains of the NE. (C) In early zygotene, consolidation of SUN-associated telomere attachment sites leads to the predominant nucleolar-side bouquet configuration (88%) characteristic of Arabidopsis meiocytes. In rare cases (12%), when SUN1/2-enriched domains occupy the opposite nuclear hemisphere, telomeres cluster accordingly, resulting in an alternative bouquet configuration. A single ASY1 focus remains associated with the 35S rDNA locus (yellow/magenta) throughout zygotene and pachytene, reflecting the persistent unsynapsed state of the NOR region. (D) As bouquet formation progresses, clustered centromeres redistribute toward the hemisphere opposite the telomere cluster, establishing the characteristic centromere–telomere opposition. Across all stages shown, the NOR remains the only chromosomal domain with a stable nuclear position, serving as a reference point for interpreting the emerging spatial organization.

Taken together, these observations suggest that early meiotic polarity in Arabidopsis arises from the interplay of three structural features: (i) the positional stability of the NOR–nucleolus complex, (ii) the heterogeneous distribution of SUN1/2 microdomains across the nuclear envelope, and (iii) the directional migration of telomeres from the nucleolus toward SUN-competent membrane regions. Although further functional analyses are needed to determine how this architecture influences pairing efficiency or recombination outcomes, the integrated spatial view provided here establishes a structural basis for interpreting early meiotic nuclear organization in Arabidopsis meiocytes.

## EXPERIMENTAL PROCEDURES

### Experimental plant material

*Arabidopsis thaliana* (L.) Heynh (ecotype Wassilewskija, WS-1) plants were grown from seeds. Plants were cultivated in a growth chamber under the following standardized conditions: day/night temperatures of 22°/19 °C; 16 h/8 h light/dark cycle; and light intensity of 150 µmol m^-2^ s^-1^.

### Preparation of male meiocytes for 3D analysis using polyacrylamide embedding

To preserve the three-dimensional (3D) architecture of male meiocytes, samples were embedded in polyacrylamide following the protocol of Hurel et al. (2018) with minor modifications. Fresh Arabidopsis floral buds (0.3-0.5 mm) were dissected by removing sepals and petals to expose the anthers. The isolated anthers and pistils were fixed in 2% formaldehyde in 1× PBS for 30 min and subsequently washed twice in Buffer A (80 mM KCl, 20 mM NaCl, 15 mM PIPES-NaOH, 0.5 mM EGTA, 2 mM EDTA, 80 mM sorbitol, 1 mM DTT, 0.15 mM spermine, 0.5 mM spermidine). 12 µL of Buffer A were placed on a silane-prep slide, and three to four buds were transferred into the drop. Anthers were gently dissected in the buffer using a needle. A 5 µL aliquot of activated polyacrylamide solution (30% 29:1 acrylamide:bisacrylamide stock supplemented with 1.25 µL of 20% ammonium persulfate and 1.25 µL of 20% sodium sulfite per 25 µL of stock) was then mixed with the Buffer A containing the anthers. A 24 × 24 mm coverslip was placed over the drop at a 45° angle, and gentle pressure was applied with forceps or a needle to release meiocytes from the embedded anthers. The polyacrylamide gel was allowed to polymerize for at least 1 h. The coverslip was then carefully removed using a razor blade. Meiocytes (chromosomes) were counterstained with DAPI (2 µg/mL) in Vectashield, and a 24 × 32 mm coverslip was mounted on top for imaging.

### Immunolabeling

Immunolabeling of selected slides containing male meiocytes were performed according to Hurel et al. (2018). To remove residual DAPI from the initial Vectashield mounting, slides were washed twice in 1× PBS for 10 min each. Slides were then incubated in 1× PBS supplemented with 1% Triton X-100 and 1 mM EDTA for 1 h at room temperature. Polyacrylamide pads were subsequently blocked with 150 µL of blocking solution for 2 h at room temperature. The blocking solution was replaced with 100 µL of primary antibody solution, and the slides were placed in a humid chamber and incubated for at least 1 h at room temperature. Primary antibodies were used at the following dilutions: anti-ASY1, anti-ZYP1, and anti-SUN1/2 at 1:250, and anti-fibrillarin at 1:100. Slides were then incubated at 4 °C for two to three nights. After primary antibody incubation, slides were washed four times in 1× PBS containing 0.1% Triton X-100 for 30 min each. Fluorophore-conjugated secondary antibodies diluted 1:250 in blocking solution (100 µL per slide) were applied and incubated for 2 h at room temperature. Slides were again washed four times for 30 min each in 1× PBS with 0.1% Triton X-100. Finally, samples were mounted in Vectashield containing DAPI (2 µg/mL), and a 24 × 32 mm coverslip was added for imaging.

### Fluorescence *in situ* hybridization

Immunolabeled slides were washed three times in 2× SSC for 10 min each. For the *in situ* localization of NORs, the Arabidopsis thaliana BAC clone T15P10 (GenBank accession AF167571), which contains 35S rRNA gene repeats, was used. Centromeres were visualized using the 178-bp centromeric satellite probe described by Naish et al. (2021). Telomeres were detected with PCR-amplified Arabidopsis-type (TTTAGGG)n repeats (fragment sizes up to 25 kb), prepared following Ijdo et al. (1991). All DNA probes were labeled by nick translation with Cy3-, biotin-, digoxigenin-, Alexa Fluor 488-, or Alexa Fluor 594-dUTP, according to Mandakova and Lysak (2016). Labeled probes were pooled, ethanol-precipitated, desiccated, and resuspended in 20 µL of hybridization buffer (50% [v/v] formamide and 10% [v/v] dextran sulfate in 2× SSC) per slide. For FISH, 20 µL of the probe mixture was applied to each chromosome-containing preparation and immediately denatured on a hot plate at 80 °C for 4 min. Hybridization was carried out overnight at 37 °C in a humid chamber. Post-hybridization washes were performed sequentially in 0.1× SSC at 42 °C, 2× SSC at 42 °C, 2× SSC at room temperature, and 4× SSC at room temperature, each for 10 min. Immunodetection of hapten-labeled probes followed the procedure of Mandakova and Lysak (2016). Finally, preparations were counterstained with DAPI (2 µg/mL) in Vectashield and mounted with a 24 × 32 mm precision coverslip.

### Imaging and image processing

Fluorescence signals were visualized and documented using an AxioImager Z2 epifluorescence microscope equipped with an Apotome module (Zeiss). Image acquisition and deconvolution were carried out in ZEN Blue software (Zeiss). IMARIS software (Oxford Instruments) was used for contrast adjustment (Channel Adjustment), three-dimensional modeling (Surface), calculations (Vantage) and video generation (Animation).

## SUPPLEMENTAL DATA

**Supplemental Video 1**. Spatial organization of chromosomes in intact Arabidopsis male meiocytes during the G2 premeiotic interphase. The G2 premeiotic nucleus was identified based on ASY1 and ZYP1 immunolabeling (grey). Telomeres (green), centromeres (red), and 35S rDNA loci (yellow) were visualized by FISH. Three-dimensional signal rendering and projections were generated using Imaris software.

**Supplemental Video 2**. Spatial organization of chromosomes in intact Arabidopsis male meiocytes during leptotene. The leptotene nucleus was identified based on ASY1 and ZYP1 immunolabeling (grey). Telomeres (green), centromeres (red), and 35S rDNA loci (yellow) were visualized by FISH. Three-dimensional signal rendering and projections were generated using Imaris software.

**Supplemental Video 3**. Spatial organization of chromosomes in intact Arabidopsis male meiocytes during early zygotene. The early zygotene nucleus was identified based on ASY1 and ZYP1 immunolabeling (grey). Telomeres (green), centromeres (red), and 35S rDNA loci (yellow) were visualized by FISH. Three-dimensional signal rendering and projections were generated using Imaris software.

**Supplemental Video 4**. Spatial organization of chromosomes in intact Arabidopsis male meiocytes during early zygotene. The early zygotene nucleus was identified based on ASY1 and ZYP1 immunolabeling (grey). Telomeres (green), which form a characteristic bouquet configuration at the nuclear periphery, together with centromeres (red) and 35S rDNA loci (yellow), were visualized by FISH. Three-dimensional signal rendering and projections were generated using Imaris software.

**Supplemental Video 5, 6**. Spatial organization of chromosomes in intact Arabidopsis male meiocytes during the late zygotene. The late zygotene nucleus was identified based on ASY1 and ZYP1 immunolabeling (grey). Telomeres (green), which form a characteristic telomere bouquet configuration at the nuclear periphery, along with centromeres (red) and 35S rDNA loci (yellow), were visualized using FISH. Three-dimensional signal rendering and projection were generated using Imaris software.

**Supplemental Video 7**. Spatial organization of chromosomes in intact Arabidopsis male meiocytes during the pachytene. The pachytene nucleus was identified based on ASY1 and ZYP1 immunolabeling (grey). Telomeres (green), which form a characteristic telomere bouquet configuration at the nuclear periphery, along with centromeres (red) and 35S rDNA loci (yellow), were visualized using FISH. Three-dimensional signal rendering and projection were generated using Imaris software.

**Supplemental Video 8**. Spatial organization of chromosomes in intact Arabidopsis male meiocytes during the early zygotene. The early zygotene nucleus was identified based on ASY1 and ZYP1 immunolabeling (grey). Telomeres (green), centromeres (red), and 35S rDNA loci (yellow) were visualized using FISH. Three-dimensional signal rendering and projection were generated using Imaris software.

**Supplemental Video 9, 10**. Spatial organization of chromosomes in intact Arabidopsis male meiocytes during the late zygotene. The late zygotene nucleus was identified based on ASY1 and ZYP1 immunolabeling (grey). Telomeres (green), centromeres (red), and 35S rDNA loci (yellow) were visualized using FISH. Three-dimensional signal rendering and projection were generated using Imaris software.

**Supplemental Video 11**. Spatial organization of chromosomes in intact Arabidopsis male meiocytes during the pachytene. The pachytene nucleus was identified based on ASY1 and ZYP1 immunolabeling (grey). Telomeres (green), centromeres (red), and 35S rDNA loci (yellow) were visualized using FISH. Three-dimensional signal rendering and projection were generated using Imaris software.

## ACKNOWLEDGEMENTS

The work was supported by the Czech Science Foundation (project no. 25-16796S to T.M.). The Plant Sciences Core Facility of CEITEC Masaryk University is acknowledged for the technical support. The CELLIM Core Facility of CEITEC Masaryk University supported by MEYS CR (LM2023050 Czech-BioImaging) is acknowledged for their support with scientific data processing presented in this paper. We thank Aurélie Chambon for training MK in the methodology, and Mathilde Grelon for hosting MK in her laboratory, for her continuous support throughout the project, and for her critical revision and valuable advice during the preparation of the manuscript.

## AUTHOR CONTRIBUTIONS

Conceptualization: TM. Methodology: MK, TM. Investigation: MK. Writing - original draft: TM with help of MK. Writing - review & editing: TM, MK, MAL. Visualization: MK. Funding acquisition: TM.

## Notes

### Competing Interest Statement

The authors have declared no competing interest.

## REFERENCES

Armstrong SJ, Franklin FC, Jones GH (2001) Nucleolus-associated telomere clustering and pairing precede meiotic chromosome synapsis in Arabidopsis thaliana. Journal of Cell Science, 114 (23): 4207–4217. 10.1242/jcs.114.23.4207

Blokhina YP, Nguyen AD, Draper BW, Burgess SM (2019) The telomere bouquet is a hub where meiotic double-strand breaks, synapsis, and stable homolog juxtaposition are coordinated in the zebrafish, Danio rerio. PLOS Genetics 15(1.): e1007730. 10.1371/journal.pgen.1007730

Cowan CR, Carlton PM and Cande WZ (2001) The polar arrangement of telomeres in interphase and meiosis. Rabl organization and the bouquet. Plant Physiology, 125, 532–538. 10.1104/pp.125.2.532

Cromer L, Tiscareño-Andrade M, Lefranc S, Chambon A, Hurel A, Brogniez M, Guérin J, Le Masson I, Adam G, Charif D, Andrey P and Grelon M (2024) Rapid meiotic prophase chromosome movements in Arabidopsis thaliana are linked to essential reorganization at the nuclear envelope. Nature Communications, 15, 5964. 10.1038/s41467-024-50169-4

Fernández-Álvarez A (2023) Beyond tradition: exploring the non-canonical functions of telomeres in meiosis. Frontiers in Cell and Developmental Biology, 11, 1278571. 10.3389/fcell.2023.1278571

Hurel A, Phillips D, Vrielynck N, Mézard C, Grelon M and Christophorou N (2018) A cytological approach to studying meiotic recombination and chromosome dynamics in Arabidopsis thaliana male meiocytes in three dimensions. The Plant Journal, 95, 385–396. 10.1111/tpj.13942

Ijdo JW, Wells RA, Baldini A, Reeders ST (1991) Improved telomere detection using a telomere repeat probe (TTAGGG)n generated by PCR. Nucleic Acids Research, 19: 4780. 10.1093/nar/19.17.4780

Mandakova T and Lysak MA (2016) Painting of Arabidopsis chromosomes with chromosome-specific BAC Clones. Current Protocols in Plant Biology, 1:359–371. 10.1002/cppb.20022

Naish M, Alonge M, Wlodzimierz P, Tock AJ, Abramson BW, Schmücker A, Mandáková T, Jamge B, Lambing C, Kuo P, Yelina N, Hartwick N, Colt K, Smith LM, Ton J, Kakutani T, Martienssen RA, Schneeberger K, Lysak MA, Berger F, Bousios A, Michael TP, Schatz MC, Henderson IR (2021) The genetic and epigenetic landscape of the Arabidopsis centromeres. Science, 374(6569):eabi7489. 10.1126/science.abi7489

Prusicki MA, Keizer EM, van Rosmalen RP, Komaki S, Seifert F, Müller K, Wijnker E, Fleck C, Schnittger A (2019) Live cell imaging of meiosis in Arabidopsis thaliana. Elife, 8:e42834. 10.7554/eLife.42834

Sheehan MJ and Pawlowski WP (2009) Live imaging of rapid chromosome movements in meiotic prophase I in maize. Proceedings of the National Academy of Sciences of the USA, 106, 20989–20994. 10.1073/pnas.0906498106

Sims J, Copenhaver GP and Schlögelhofer P (2019) Meiotic DNA repair in the nucleolus employs a non-homologous end-joining mechanism. The Plant Cell, 31, 2259–2275. 10.1105/tpc.19.00367

Sims J, Rabanal FA, Elgert C, von Haeseler A and Schlögelhofer P (2021) It Is Just a Matter of Time: Balancing homologous recombination and non-homologous end joining at the rDNA locus during meiosis. Frontiers in Plant Science, 12, 773052. 10.3389/fpls.2021.773052

Tomita K and Cooper JP (2007) The telomere bouquet controls the meiotic spindle. Cell, 130, 113–126. 10.1016/j.cell.2007.05.021

Varas J, Graumann K, Osman K, Pradillo M, Evans DE, Santos JL and Armstrong SJ (2015) Absence of SUN1 and SUN2 proteins in Arabidopsis thaliana leads to a delay in meiotic progression and defects in synapsis and recombination. The Plant Journal, 81: 329–346. 10.1111/tpj.12730

Zeng X, Li K, Yuan R, Gao H, Luo J, Liu F, Wu Y, Wu G and Yan X (2018) Nuclear envelope-associated chromosome dynamics during meiotic prophase I. Frontiers in Cell and Developmental Biology, 5, 121. 10.3389/fcell.2017.00121

Zickler D and Kleckner N (1998) The leptotene–zygotene transition of meiosis. Annual Review of Genetics, 32, 619–697. 10.1146/annurev.genet.32.1.619

